# Signaling gradients in surface dynamics as basis for planarian regeneration

**DOI:** 10.1101/733246

**Authors:** Arnd Scheel, Angela Stevens, Christoph Tenbrock

## Abstract

We introduce and analyze a mathematical model for the regeneration of planarian flatworms. This system of differential equations incorporates dynamics of head and tail cells which express positional control genes that in turn translate into localized signals that guide stem cell differentiation. Orientation and positional information is encoded in the dynamics of a long range *wnt*-related signaling gradient. We motivate our model in relation to experimental data and demonstrate how it correctly reproduces cut and graft experiments. In particular, our system improves on previous models by preserving polarity in regeneration, over orders of magnitude in body size during cutting experiments and growth phases. Our model relies on tristability in cell density dynamics, between head, trunk, and tail. In addition, key to polarity preservation in regeneration, our system includes sensitivity of cell differentiation to gradients of *wnt*-related signals relative to the tissue surface. This process is particularly relevant in a small tissue layer close to wounds during their healing, and modeled here in a robust fashion through dynamic boundary conditions.

## 1 Introduction

Planarians are nonparasitic flatworms commonly found in freshwater streams and ponds [62, 64] with a body size in the *mm*-scale. They possess the ability to regenerate after rather severe injuries to their body. In extreme cases, the entire organism can regenerate perfectly from a fragment just 0.5% of the original size. The potential of the relevant processes has helped direct a tremendous amount of attention toward studying mechanisms during regeneration in these organisms.

The purpose of this paper is to present a minimal model, informed by experimental data, that reproduces this fascinating regenerative behavior. To the best of our knowledge, the mathematical model we introduce here, is to date the only one, based on reacting and diffusing species, that is able to correctly recover most of the typical cutting and grafting experiments. In particular, our model shows preservation of polarity during regeneration small tissue fragments. This reflects experimental findings where small tissue parts cut from a planarian regenerate to a fully functioning and intact organism with head and tailpositioned such, that the original orientation of the tissue fragment is respected. On the other hand, when tissue parts are cut from a donor and implanted into a host, the newly created planarian integrates the positional information of the tissue fragment from the donor with the new positional information it obtains from the host.

The mechanism proposed here resolves a conundrum in modeling efforts. In fact, many models of spontaneous formation of finite-size structure in unstructured tissue allude to a Turing type mechanism to select a finite wavelength. These types of models have often been discussed in the context of regeneration phenomena in hydra, which exhibit similar robustness features as planarian regeneration that we focus on here. Regeneration of hydra has been one of the central motivations for many studies of activator-inhibitor type systems [40]. Such Turing mechanisms however do not scale across several orders of magnitude as seen in the patterning of planarians, nor do they incorporate robust selection of polarity. In the remainder of this introduction, we briefly describe the species and kinetics in our model, the role of dynamic boundary conditions, and the key experiments that we aim to mimic in our modeling efforts.

### Cell types and signaling agents

We focus on the ante-posterior axis, describing dynamics in a one-dimensional domain *x* ∈ [−*L, L*], populated by different cell types and signaling agents. Head cells *h* and tail cells *d* are generated from stem cells *s* through differentiation. Head and tail cells generate signals *u_h_* and *u_d_*, respectively. In addition, we model production and diffusion of a long-range *wnt*-related signal *w*, in the following often referred to as *wnt*-signal in short, that encodes orientational information through its gradient. We model random motion of cells and diffusion of signal molecules *U*(*t,x*) = (*s,h,d,u_h_,u_d_,w*)(*t,x*) through diffusion coefficients *D_j_*, and postulate kinetics

- *F_s_*(*s, u_h_, u_d_*) for proliferation and differentiation of stem cells;
- *F_h_*(*s, h, u_h_*), *F_d_*(*s, d, u*_d_) for differentiation of stem cells into head and tail cells and death of those;
- *F_u_h__*(*h, d, u_h_*), *F_u_d__*(*h, d, u_d_*) for regulation of the signals *u_h_* and *u_d_* by head and tail cells;
- *F_w_*(*h, d, w*) for degradation and production of the *wnt*-related signal by head and tail cells, respectively.

The kinetics of *F_s_* regulate a near-constant supply of stem-cells. The dynamics for the signals *u_h_* and *u_d_*, which are produced by head and tail cells but degraded by reactions with tail and head cells, respectively, encode tristability between a head-only, a tail-only, and a zero state. The long-range signal *w* is produced by tail cells and degraded by reactions with head cells. As a result, a healthy planarian consists of a high concentration of *h* near *x* = −*L*, a high concentration of *d* near *x* = *L*, and a near-constant gradient of *w* in *x* ∈ (−*L, L*), see Figure 2.1.

**Figure 2.1:**
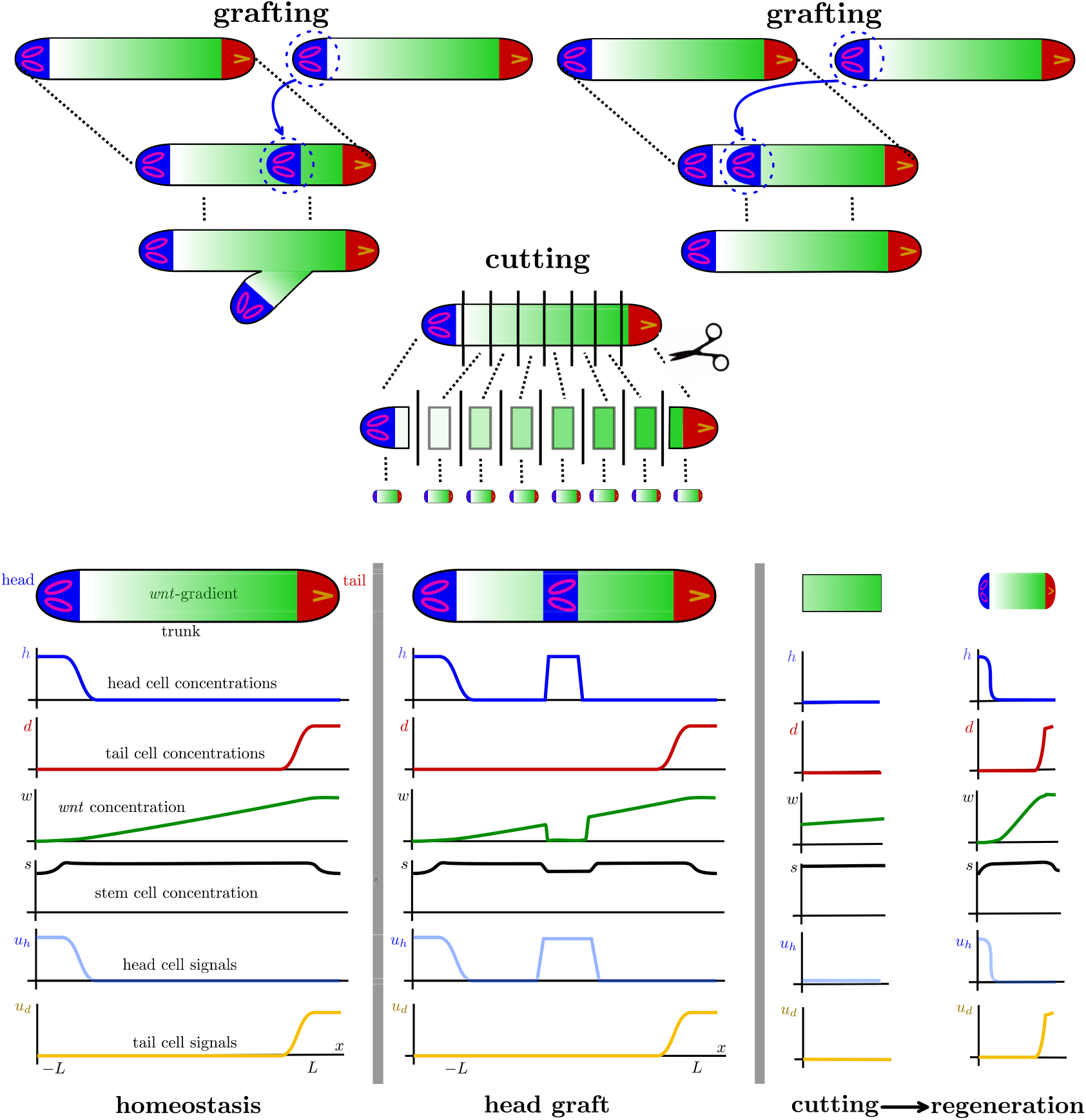
Typical experiments and their representations in terms of our mathematical model. Top left: Schematic illustration of head grafting and regeneration of a two-headed animal. Top right: Head grafting and regeneration into a normal planarian with one head and one tail. Middle row: Cutting experiment and regeneration of 8 planarians. Bottom row: Schematic illustration of experiments on planarians and associated spatial distribution of concentrations *h, d, w, s, u_h_, u_d_*. Homeostasis (left) has head cells concentrated on the left, tail cells concentrated on the right, a *wnt*-gradient directed towards *wnt*-production in the tail, roughly constant stem cell populations with slightly decreased populations where differentiation into head and tail cells occurs, and head and tail signals are closely mimicking the distribution of head and tail cells. In grafting experiments (center), the head region of a donor planarian is grafted into a host planarian, retaining roughly the distribution of cell concentrations and signaling molecules from its original location. In cutting experiments (right), a thin fragment (here from the trunk region) retains a small *wnt*-gradient but no head or tail cells. Regeneration in this context refers to reestablishing head and tail cell populations while preserving polarity.

### *wnt*-related regulation of cell types

We emphasize here that differentiation rates do *not* depend on *w* in our model. In other words, we do not incorporate mechanisms that convert levels of the *wnt*-related signal (or other long-range signals) into positional information that in turn directs the differentiation process. Such mechanisms, often referred to as the French-flag model [86, 87], are widely assumed to play a key role in early development and likely also have significant influence on the regeneration of planarians [64]. Here, we omit this dependence for several reasons. First, we believe that such positional information is inherently inapt at explaining the preservation of polarity, since the absolute levels of the *wnt*-signal play apparently little role in the regeneration from small tissue fragments. This is exemplified by the robustness of regeneration and polarity in regards to the location of the cut out tissue fragment, near head, near tail, or from central parts of the body. Second, such information is not necessary in the early stages of the regeneration, but seems to be relevant only later, when also the size of functional regions such as head or tail are regulated. Lastly, related to this latter point, there is little concrete information so far on the nature of biological mechanisms that would accomplish this translation of, for instance, *wnt*-signal levels into an information for differentiation of stem cells in a robust fashion. Therefore we see value in not simplifying the modeling task by the addition of such somewhat poorly substantiated regulatory mechanisms.

Rather than using positional information based on the absolute value of the *wnt*-signal, our mathematical model is crucially based on the detection of gradients of *wnt*, in particular the orientation of the gradient relative to the respective body edge (encoded in the outer normal of the boundary). We see this orientation of the gradient as an in some sense necessary, minimal information, to guide regeneration while at the same time preserving polarity. This detection of gradient orientation is encoded in boundary dynamics that we shall explain next.

### Boundary dynamics

Cutting experiments eliminate head and/or tail cells, therefore ultimately influence the *wnt*-pathway and destroy the associated signaling gradient. We therefore include boundary dynamics into our model which represent wound healing processes at wound edges that in particular lead to regeneration of head, tail, and ultimately reestablish the signaling gradient of *w*. This is due to fast differentiation of stem cells close to the wound edges. Differentiation into head versus tail cells is guided by information on the normal derivative of the residual signal *w* in the body fragment. We model these boundary dynamics through dynamic boundary conditions (sometimes referred to as Wentzell boundary conditions in the literature). Therefore, we introduce values *U*_±_(*t*) of all cell and signal concentrations at the left and right boundary compartment, respectively, that specifically act as Dirichlet conditions for the chemical species *U*(*t,x*). We then introduce differential equations for *U*_±_(*t*) based on a diffusive flux *∂_ν_ U* to ensure mass balance, the *same* kinetics *F_j_*, now evaluated at the boundary, and, crucially, one additional term that accounts for strong differentiation of stem cells into head and tail cells, namely *s* · Ψ_*h*/*d*_, during wound healing. We postulate that, as a key ingredient to robust regeneration and preservation of polarity, the production rate Ψ_*h*/*d*_ = Ψ_*h*/*d*_(*h, d, ∂_ν_w*) depends sensitively on the normal derivative of the long-range signal *w*, measuring roughly the sign of *∂_ν_w*. We model this switching behavior through a rate function

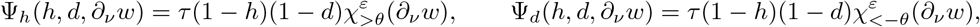

where *τ* ≫ 1 is the rate and 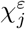 are smooth versions of the characteristic function such as

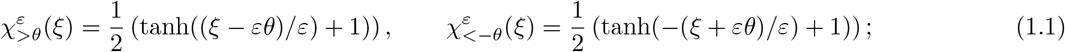

see Figure 3.1 for an illustration. The steepness *ε*^−1^ of the smoothed characteristic function can be interpreted as a sensitivity of the production in regards to small gradients. One can envision many scenarios that enable stem cell differentiation to be guided by gradients of a chemical signal, for instance through comparing signal strength spatially or temporally. This latter process can be enhanced by directed motion of stem cells or progenitors into the respective directions. Indeed, migratory stages of progenitors from their place of birth to their site of terminal differentiation are discussed in [64]

**Figure 3.1:**
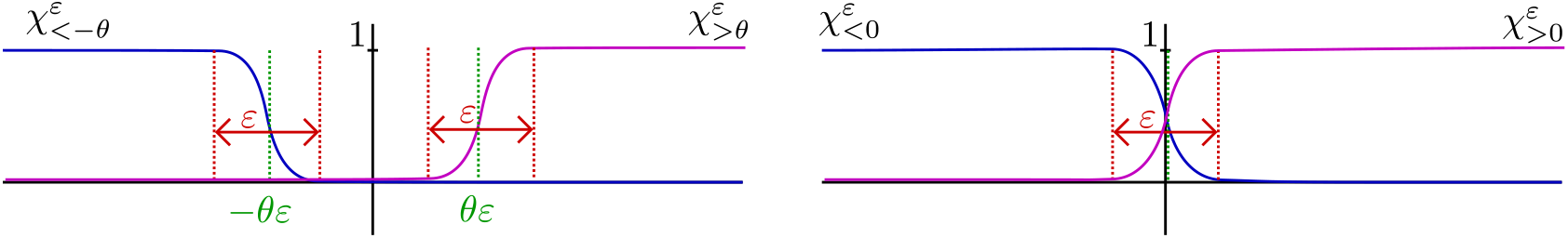
Schematic plot of the indicator functions *χ* that detect positive and negative values of the gradient, respectively, with offset *θ* and sensing thresholds *ε*.

From a mathematical perspective, it is interesting to note that dynamic boundary conditions of this type cannot be readily replaced by, say, Robin boundary conditions, by letting for instance rates of boundary dynamics tend to infinity. This mathematical curiosity bears consequences on modeling assumptions, implying for instance the presence of a distinguished boundary/body region. Such a region has recently also been discussed in the experimental literature [64], where the concept of poles, separating head regions from trunk, is attributed a key role in regeneration. On the mathematical side, we make this curious role of dynamic boundary conditions precise in a simple reduced model toward the end of this paper.

### Regeneration from cutting, grafting, and growth — simulations

The model ingredients presented thus far can, to a large extent, be motivated through experimental data. We validate the modeling effort by displaying results from numerical simulations that mimic basic experiments on regeneration, such as cutting, grafting, and growth; see Section 3.

### Regeneration — analysis

We provide some analytical understanding of the key ingredients of our mathematical model by deriving a reduced system, consisting of an order parameter *c* that lumps concentrations of head and tail cells into a scalar quantity, coupled to the long-range *wnt*-related signal *w*. In this reduction, we clearly illustrate how the nonlinear boundary fluxes restoring head and tail cell concentrations, dependent on the information of normal derivatives of the gradient, initiate and organize the regeneration process. We outline a (in)stability analysis that points to the origin of regeneration and we pinpoint failure of this process when dynamic boundary conditions are relaxed to, say, nonlinear mixed (or Robin) boundary conditions. Lastly, we distill the key feature, restoration of a signal gradient through boundary, respectively body edge, sensing, into a dramatically oversimplified scalar model for the *wnt*-related signal. In this scalar model, we then clearly outline the limits of regeneration and point to curious oscillations, caused by coupling of scalar, non-oscillatory dynamics in the boundary to the diffusive signal field in the bulk and the resulting delayed feedback mechanism.

### Outline

We review biological experiments that motivate our model and compare with other systems in the literature in Section 2. We then introduce our mathematical model and describe numerical simulations that mimic planarian regeneration in various scenarios in Section 3. Section 4 contains analytical results that reduce dynamics to a two-species and ultimately a scalar reaction-diffusion system, thus exhibiting the key dynamic ingredients of our original model. We also discuss the necessity of the somewhat non-standard boundary conditions and the limitations of the tristability mechanism. We conclude with a brief summary and discussion.

## 2 Planarian regeneration — experiments and models

We give a brief overview of planarians in Section 2.1, describe experiments with a short comparison to our findings from the mathematical model in Section 2.2, give genetic information in Section 2.3, and summarize the existing mathematical literature in Section 2.4.

### 2.1 Basics of planarian regeneration

In planarian pluripotent adult stem cells (neoblasts), are the source of all cell types. Neoblasts divide, differentiate, and differentiated cells die after some time. There is a continuous turnover of cells, still the flatworm maintains cell type proportions. Understanding the mechanisms responsible for this dynamic steady state is one important objective in research on planarians.

Planarians exhibit a bilateral symmetry, three distinct body axes, a well-differentiated nervous system including a brain, a gastrovascular tract, a body-wall musculature, and they consist of three tissue layers [32]. The best experimental data is available for the species “Schmidtea mediterranea”, which is 1 *mm* to 20 *mm* long [62] and consists of 100.000 to more than 2.000.000 cells. At least 20 to 30 different types of differentiated cells are observed. The broadly distributed pluripotent population of neoblasts constitutes possibly up to 30 % of the total cell population [7] and are the only proliferating cells in planarians [75].

Regeneration in our context is the ability of an organism to replace lost or damaged tissue. The regenerative abilities of humans are limited to special tissues only. Examples for regeneration after injury in animals include regeneration of deer antlers, fins of fish, tails of geckos, or complete limbs in some crabs or salamanders. Planarians and hydra are among the few species that seem to possess a nearly unlimited regenerative ability. They recover from practically every injury, and regenerate when aging. A small fragment cut out of a planarian, as small as about 0.5%, can regenerate a full animal, including in particular an intact brain [62]. Some asexual planarian species even reproduce by tearing themselves apart and subsequently develop into two intact worms [32]. Clearly, the study of this behavior holds tremendous appeal when considering the potential of a better understanding of regeneration of tissues in the human body, for instance parts of the heart muscle after an infarct.

### 2.2 Experiments

We give a short overview of classical experiments and refer to [8, 10, 6, 11, 13, 16, 66, 42, 57, 58, 67, 72, 47, 43, 41, 65] for more details. We also briefly point to and summarize our findings in relation to these experiments.

#### 2.2.1 Cutting experiments

Cutting tissue off a planarian results in regeneration of both parts into an intact organism, more or less independent of position, size, or direction of the cut, with few exceptions, discussed below. Here we focus on regeneration of the anterior-posterior (AP, or head-to-tail) axis, and think mostly of dissections transverse to the AP axis into strips whose width is a fraction of the original length of the planarian body; see Figure 2.1. Each of these tissue strips will regenerate into a complete animal after several weeks. In tissue pieces cut close to head or tail, regeneration of a new head might occur in as little as three to four weeks. Complete regeneration, including restoration of the right proportions, usually requires two to three months, depending on the shape of the respective tissue fragment [41]; see Figure 2.1 for schematics of experiments and representation in our model. Polarity in such fragments is preserved, that is, a head will regenerate at the edge that has been closer to the head before, and the tail will regenerate at the opposite end. In particular, neighboring cells in a planarian will regenerate as either head or as tail after they have been separated by a cut! Among the few exceptions are very long or very short tissue fragments, which sometimes regenerate a second head instead of a tail. Moreover, tissue parts that are too small or too thin are not able to regenerate at all.

Our mathematical model reproduces all of these experiments robustly, for a wide range of parameter values, with a few exceptions that we note below; see Figure 3.4 for simulations of our full system and Figures 4.2, 4.3, and 4.4, 4.5 for simulations of a reduced model and for a scalar equation. In our mathematical model head and tail cells are treated equivalently with equal weights to cells and corresponding reaction rates. As a consequence, simulations are not able to produce a bias between two-headed and two-tailed animals. Specific outcomes depend only on small fluctuations in initial conditions when recovery is not robust, that is for very small or or for large fragments. Matching experiments, we found recovery somewhat more robust for tissue fragments from the center of the body when compared to equally sized fragments cut from regions near head or near tail. In the analytically best understood case of the scalar model, we observed that when changing parameters toward a regime where regeneration is impossible, oscillations occure in the recovery process before, upon further change of parameter values, the *wnt*-gradient fails to recover at all. These are the exceptions from the otherwise matching behavior of our mathematical model in comparison to experimental findings.

##### Grafting experiments

In grafting experiments a tissue fragment of an intact donor planarian is cut out and transplanted into another healthy host planarian, whose regeneration is then studied. The donor usually regenerates as described before, whereas the host may develop new phenotypes; see again Figure 2.1. Most interesting are experiments where parts of the head of the donor are transplanted into different positions within the host. Depending on size and position of the donor head tissue within the host, the host will either regenerate into a completely normal animal, with the grafted head vanishing, or it will generate a second axis at the position of the transplant such that the AP axis splits into two branches of equal length exhibiting two heads and one tail, or it will generate a two-headed planarian. Data suggests that outgrowth is more likely if there is a larger distance between the head transplant and the head of the host. Similarly, outgrowth seems to become more likely for larger head transplants and for tissue taken closer to the head of the donor.

Transplanting a donor tail into the upper part of the host might reorganize the host’s middle region, sometimes regenerating a second pharynx in opposite direction, possibly also leading to outgrowth. Transplanting a complete lower donor fragment below an upper fragment of a bisected host, such that just an intermediate strip for a complete animal is missing, will always regenerate the intermediate strip.

We mimic such experiments in simulations of our mathematical model; see Figure 3.5 and 3.6 for the full system and Figure 4.2 for the reduced model and confirm to some extent the observed dichotomy between the emergence of a new head and the vanishing of the grafted head during regeneration. As a caveat, the insight into these phenomena from our model is limited by the fact that we do not model regulation of organ growth through chemical signals.

##### Growth and Shrinking

Adult planarians of the species “Schmidtea mediterranea” are roughly 20 *mm* long if well fed, and can shrink to about 1 *mm* if starved, and regrow when food supply is restored, keeping relative proportions and ratios of cell populations intact [7]. During regeneration after cutting experiments, this ability is crucial: if, for instance, a tissue fragment is cut from close to head or tail, it will initially lack a pharynx, and therefore draw resources from the fragment shrinking to as little as a tenth of the original fragment size.

We show experiments that show robust behavior over a wide range of body sizes and growth rates; see Figure 3.7.

##### Hydra

The regeneration of hydra, a freshwater polyp, shows similarities to that of planarians, with comparable outcomes for most of the experiments mentioned above; see [80, 33, 71, 70, 12, 84] and references therein. Different from planarians, hydra stem cells are distributed exclusively inside the body region, below the epithelium, such that a fragment consisting only of head or foot cells will not regenerate; see [1] for one of the very few quantitative studies on induction of additional foot or tail axes.

An interesting *dissociation experiment* for hydra has not been described for planarians so far. When a tissue fragment of hydra is pressed through a net and the resulting small fragments are reorganized randomly, a bulb of hydra tissue without any sense of polarity and positional information arises. This tissue fragment will subsequently regenerate head and foot structures. However, depending on the number of involved cells, it might generate several heads and body axes that will separate only later [48]; compare Figure 3.6 for a related set-up in our model.

### 2.3 Genetic information

In order to identify gene expression and their spatial and temporal dynamics in planarians, the amount of RNA produced from specific genes during protein synthesis is measured, for instance through in situ hybridization or northern blot [49]. Therefore genes can be identified, which are mainly expressed close to the head or tail, or after wounding and feeding. To artificially influence gene expression by RNA interference, targeted mRNA molecules are neutralized, thus synthesis of messenger molecules stops at an earlier point in time. With this manipulation it can be tested, e.g. which of the genes being expressed within the head region are actually necessary to regenerate a head.

So far, in both hydra and planarians, it does not seem possible to track the corresponding messenger molecules directly. Corresponding antibodies are not yet available. The experimental methods described above, do allow however, to some extent, for an analysis of the production dynamics and functioning of these signals. The identification of their final distribution in the planarian body seems out of reach at the moment, but would be important to know for further mathematical modeling efforts. We refer to [51], [64] for a review on biological results pertinent to our mathematical model presented here.

#### Gene expression sites

The three main body axes in planarians are organized by different signaling systems. Knock-out experiments suggest that these systems act quite independently [61]. We therefore focus on genes which organize the anterior-posterior (AP) axis. Information about the other axes, and more details on the AP axis can be found in [2, 61, 52, 22, 4, 19, 63].

Genes which are expressed close to the head are *notum, sFRP-1, ndl-4, prep*, and *sFRP-2*, among others. While being expressed mainly in the upper half of planarians, some of these genes are also expressed at the lower tip of the pharynx region in the center of the body. Genes which are expressed closer to the tail include *wnt1, fz4, wnt11-2, wnt11-1, wnt11-5* (or *wntP-2*), and *Plox4-Dj*. Among the head related genes, *notum* and *sFRP-1* are expressed more locally, while expression of *sFRP-2* extends from the planarian head tip to its center. Similarly, among the tail related genes, *wnt1* is expressed very locally, while *wnt11-5* is expressed from the tail tip to the center region in a graded fashion. Local expression here refers to only a few cells at the very tips of the planarian that are responsible for the production of associated molecules, most likely subepidermal muscle cells [85].

The time dynamics of gene expression after wounding are described in [4, 53, 81, 54]. When a tissue strip of a planarian is cut from the center of the body, lacking head and tail, one can distinguish between genes that are activated early, during the first 0 – 18 hours after amputation, or later. Further one can identify genes that are expressed asymmetrically, either at head or at tail wounds but not at both. Genes, that are expressed very early after wounding include *wnt1, wnt11-5, notum, sFRP-1*, but only *notum* is expressed asymmetrically, that is, at wounds that face missing head structures. Genes such as *wnt1* are expressed first at all wound sites and their expression is normalized to the behavior we described before, only later. Among 128 wound induced genes only *notum* shows the polarized expression [88], mentioned already, and only the downstream factor of *wnt*-signaling, *β-catenin*, seems to influence this asymmetric expression [68].

#### Knock-out experiments

While it is highly desirable to acquire information about the action of genes and especially their corresponding messenger molecules, a clear picture seems elusive at this point, due to the shear number of genes involved in regeneration processes; see [61] and [51] for recent reviews.

Inhibition of most genes that are expressed early after wounding with expression patterns related to head or tail identities causes regeneration failures of the respective body region at their sites of expression. For instance, inhibition of *notum* and head amputation prevents head regeneration, while inhibition of *wnt1* and tail amputation prevents tail regeneration. In these cases the planarian will regenerate with two tails or two heads, respectively. Considering *wnt1* and *notum* as one of the first expressed genes after wounding it is interesting to note that they act in an antagonizing manner [27].

Most important for establishing and maintaining the AP axis/polarity appears to be the canonical *wnt*-pathway. This can be demonstrated by influencing the level of *β-catenin* inside the cell. High levels of *wnt* signaling correlate with high levels of *β*-catenin within the cytoplasm. The inhibition of *β*-catenin itself leads to different phenotypes [2]. If low doses of inhibitor *dsRNA* are injected into a planarian and the tail is removed, the wound closes but no tail regenerates. On the other hand, higher doses will lead to regeneration of a second head at the tail wound with a second opposing pharynx in the middle of the planarian body. If the doses are further increased, the pharynx (both) become disorganized and ectopic eyes appear. A complete inhibition of *β-catenin* leads to a radially shaped planarian which has head-related structures (nerve cells, eyes, etc.) at every side, even without tail amputation. In this context, we remark that more recently, an organizing *β-catenin* concentration gradient, with maximum at the head and minimum at the tail has been confirmed experimentally [74, 73].

#### Gene expression in hydra

The hydra homologue of the above mentioned genes *wnt, dishevelled, gsk3, tcf* and *β-catenin* seem to act in a comparable way. Studies on expression pattern and knock-out experiments can be found in [56, 25, 55, 28, 20]. The similarities appear strong enough to suggest that models developed here for planarians would carry implications also for hydra. More generally, the *wnt*-pathway appears to be more widely conserved during evolution in many species, beyond hydra and planaria.

### 2.4 Mathematical models in the literature

Efforts toward mathematical modeling the emergence of pattern or structure in seemingly unpatterned developmental stages of organisms go back to Alan Turing’s seminal work [77], where he studied the possibility of pattern formation in systems of reaction-diffusion equations, based on disparate diffusion length scales. He included in particular the case of vanishing diffusion and elaborated on the selection of long-, finite-, and short-wavelength patterns, as well as oscillatory, traveling-wave patterns, depending on reaction constants. The key observation was that it is possible that the spatially extended system can be unstable, even when simple kinetics do not exhibit instability, hence the terminology of diffusion-driven instabilities and pattern formation. He also modeled the development of tentacles in hydra [77] but could not finish a follow-up manuscript on mathematical modeling of developmental processes because of his untimely death. The system of two reaction-diffusion equations from Turing’s paper has been tremendously influential in the literature and was used and built upon by others to model, for instance regeneration in hydra, in particular in [21] where the notion of activator-inhibitor system was introduced. For other models based on such kind of dynamics see [45], [46] and the references therein. Turing’s mechanism has also been incorporated into descriptions and models of regeneration after cutting experiments in hydra and planarians and a certain class of grafting experiments. However, the main feature of Turing’s mechanism, the selection of a preferred finite wavelength, like for the tentacles of hydra, there turns into a drawback since patterns in such models strongly depend on the domain size, exhibiting more concentration maxima and minima on larger domains, while the pattern in the regeneration of the planarian body axis develops and is robust over several orders of magnitudes of variation in body size. Attempts in the literature to address this conundrum are typically built on additional species that evolve dynamically and affect reaction constants in Turing’s mechanism in ways that change Turing’s selected wave length. In well designed contexts, one can thus achieve scaling invariance of patterns over large changes in domain size [50, 79, 78, 83], thus providing mechanisms and models for experimental evidence of scale-independent patterning as for instance in zebra fish [3].

With a focus back to planarians, regeneration of a pattern would not necessitate the formation of multi-modal structures but rather necessitate establishing a monotone signaling gradient. Generation of such monotone structures has in turn been studied mathematically quite extensively in other modeling contexts, for instance in phase separation [18] and in cell polarization [44]. In the reaction-diffusion context, the simplest formulations lead to 2-species mass-conserving reaction-diffusion systems, which can exhibit pattern formation, albeit with a wavelength proportional to the domain. Patterns of smaller wavelength are unstable against coarsening although coarsening can be slow or even arrested for small or vanishing diffusivities.

Motivated by the receptor-ligand binding model [69], a more complex model for regeneration in hydra was developed in [34], again based on diffusion driven instabilities with features of activator-inhibitor models. Later, introduction of hysteresis into such models allowed the authors to correctly recover grafting experiments in numerical simulations of this system of six coupled ordinary and partial differential equations for most relevant scenarios [35]. Intuitively, mechanisms based on hysteresis or multistability are well suited to describe the outcome of grafting experiments as higher values of suitable variables, which correspond to head identities, are stabilized, independent of position or domain size. Additionally, effects such as the absorption of very small transplants might be explainable, too. In fact, our model contains some features of the associated multistability. On the other hand these models typically do not generate relevant patterns robustly in situations corresponding to cutting and dissociation experiments. Systems of ordinary differential equations (vanishing diffusivities) coupled to partial differential equations have been studied extensively, yet more recently in regard to their pattern-forming capabilities [37, 36, 23, 38, 31, 39], finding for instance stable patterns and unbounded solutions developing spikes. In a different direction, the role of hysteresis in diffusion-driven instabilities and de novo formation of stable patterns was investigated in [24, 30]

Lastly, we point out that our analysis connects with recent efforts to model and understand the role of distinguished surface reactions and bulk-to-surface coupling in morphogenesis [17, 15, 29, 59, 60]. Envisioning for instance two species reacting and diffusing with equal diffusion constant on a surface, but one of the species diffusing rapidly into, through, and back out of the bulk, one immediately finds the disparate effective diffusivities necessary for pattern formation in Turing’s mechanism, thus providing a biologically realistic mechanism for robust pattern formation.

We refer to [76] for a more in-depth discussion of recent models based on diffusion-driven pattern formation, and in particular also to the modeling and simulation of some variants of the mathematical model discussed here.

As we shall see below, our model bears little resemblance with the modeling efforts described in this section. It is arguably simpler, at least in the reduced forms of Section 4, than most models based on diffusion-driven instabilities, and possibly more versatile in the experimental phenomenology that can be robustly reproduced.

## 3 Cell types, signals, and boundary dynamics

We introduce cell and signal densities motivated by the discussion in Section 2, together with the associated production rates in Section 3.1. We then discuss dynamic boundary conditions and the additional gradient-sensing effect in Section 3.2. Section 3.3 contains numerical results on cutting, grafting, and growth, illustrating the features of our mathematical model.

Much of the mathematical formulation, including cell types and signals, is based on the thesis of one of the authors [76], but we deviate in our treatment of boundary conditions and gradient sensing.

### 3.1 Cell types, signals, and rates in the bulk

We restrict ourselves to processes regulating the anterior-posterior (AP) axis and therefore consider a simple onedimensional spatial domain *x* ∈ [−*L, L*].

#### Cell types

We model three cell types depending on time *t* and position *x* within the planarian,

|  |  |
| --- | --- |
| $s(t, x)$ | density of stem cells |
| $h(t, x)$ | density of head-specific cells |
| $d(t, x)$ | density of tail-specific cells |

#### Signals

Next, we distinguish between signals, whose actions are strongly associated with one of the cell types and are typically very localized, and signals with long-range effects that are influenced by more than one cell type. Short-range signals in our model, acting locally, are

|  |  |
| --- | --- |
| $u_h(t, x)$ | density of signals related to head-specific cells |
| $u_d(t, x)$ | density of signals related to tail-specific cells |

In addition, we consider a longer range *wnt*-related signal that establishes a full body gradient. It is produced by tail cells and degraded by reactions with head cells,

|  |  |
| --- | --- |
| $w(t, x)$ | density of lumped *wnt*-signaling |

The idea here is that body fragments retain a gradient in the concentration of this signal *w* which can then provide clues for polarization in regeneration. In fact, much of the current understanding of planarian regeneration rests on the idea of such a full body gradient of *wnt*-signaling [2], although details are not completely understood. Measurements of *β-catenin*, a cell-internal downstream factor of *wnt*-signaling [19] seem to support this idea of a long-range gradient in concentration, decreasing from tail to head [73, 74]. Expression of *wnt1* is clearly associated with tail identities [53] and leads to accumulation of *β-catenin* inside the cells [14]. On the other hand *notum*, a *wnt* antagonist [27] expressed locally at the tip of the head [61], is the only gene that shows polarized expression directly after wounding [88] and is required for head regeneration [68]. Its expression is exclusively affected by *β-catenin* signaling [68, 54]. It therefore appears that these mutual dependencies of *β-catenin* and members of the *wnt*-signaling family together with its inhibitors form a *wnt*-related signaling gradient over the full body.

Our variable *w* represents this long-range *wnt*-signaling family. Although it is often argued that a corresponding signal, produced by head cells and degraded by tail cells, play a role in regeneration, we do not incorporate an additional variable modeling such a signal since at present understanding the additional information in such a signal would be equivalent to 1 − *w* and therefore not contribute in a mathematically essential way.

#### Reaction and diffusion

We model the space-time evolution of these six concentrations *U* = (*s,h,d,u_h_,u_d_,w*) via a reaction-diffusion system

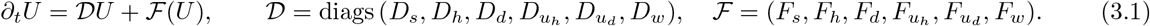

Boundary conditions are explained in Section 3.2. First, we discuss the specific forms of the production rates *F_j_*, *j* ∈ {*s, h, d, u_h_, u_d_, w*}.

#### Stem cell production and differentiation rates

We assume that stem cells proliferate, undergo apoptosis [9], and differentiate irreversibly into other cell types guided by positional control genes, that is, the head and tail related signals,

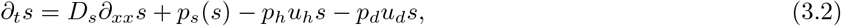

where 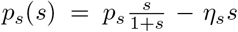 encodes saturated proliferation and apoptosis. The localized signals *u_h_,u_d_* trigger differentiation into the associated cell lines and are subsequently produced by these cells. This system acts during both, normal tissue turnover and regeneration [62].

#### Head and tail cells

Head and tail cells do not proliferate but result from differentiation of stem cells and undergo apoptosis, leading to

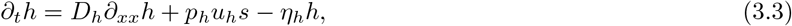

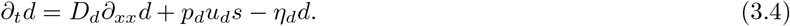

#### Signal production and depletion rates

The differentiated cells produce the signal associated with their cell type and degrade signals associated with other cell types,

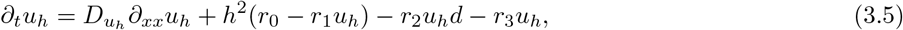

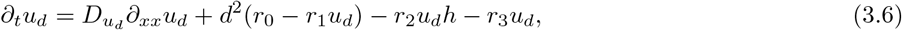

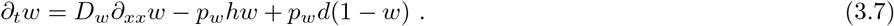

Specifically, *r*_0_ and *r*_1_ measure saturated production of cell-type related short-range signals, *r*_2_ degradation of short-range signals in reactions with the respective other cell type, and *r*_3_ otherwise induced losses of the signals. The quadratic dependence of production of *u_h_* and *u_d_* on *h* and *d*, respectively, is relevant, since replacing this reaction by first-order kinetics, that is, linear in *h, d*, would not result in tristability and spontaneous growth of head and tail regions; see (4.1) in our analysis and model reduction, Section 4.1.

The kinetics of the *wnt*-signaling complex *w* are motivated by simple degradation via head cells and saturated production via tail cells. We have used the same parameter *p_w_* for degradation and production rates. We tested differing rates with qualitatively similar outcomes. Reducing for instance to a smaller saturated production rate, the tail simply regenerates more slowly, that is, only after the head is formed. Grafting experiments seemed insensitive to changes in rates. Growth experiments fail only when the saturated production rate is too small to compensate for the effect of dilution.

### 3.2 Dynamic boundary conditions and *wnt*-signaling gradients

Tissue loss, for instance from cutting, leads to a local peak of stem cell differentiation [82]. This reaction, which is specific to wounds with loss of tissue is not completely understood [51], but appears to be important for regeneration.

We choose dynamic (or Wentzel) boundary conditions to model such strong effects near cutting sites during wound healing. These somewhat non-standard boundary conditions supplement the reaction-diffusion system (3.1) with inhomegeneous Dirichlet boundary conditions and evolution equations for the time dependent Dirichlet data,

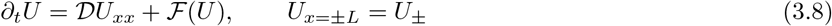

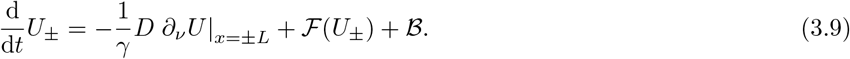

We think of the concentrations *U*_±_ as measures for spatially constant concentrations in a *boundary compartment*. Within this boundary compartment, close to the respective body edges, the same reactions 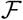 as in the central body parts/bulk take place, and diffusive flux terms 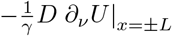 guarantee mass conservation up to production terms, as can be seen from the short calculation

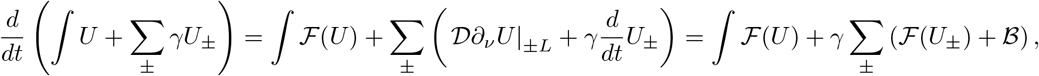

where *γ* is the size of the boundary compartment (or the body edge), measured in unit lengths. The key difference between these boundary compartments and the bulk of the domain is the presence of the terms 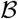 that we shall describe, below.

The main rationale for dynamic boundary conditions is the need to model dynamics in a thin but finite-size boundary compartment close to the body edges, where kinetics are slightly different from the planarian trunk. We assume that the boundary compartment is comparatively small such that concentrations of all cells and signals are constant inside, that is, *U*_±_ does not depend on space.

Dynamic boundary conditions can be compared to nonlinear fluxes. Formally, letting the size of the boundary compartments *γ* tend to 0 and assuming at the same time rapid reactions 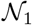 within these boundary compartments

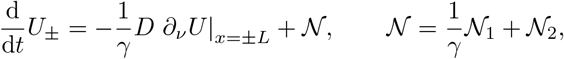

one finds in the limit *γ* = 0 the mixed boundary condition 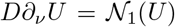. We shall see later, in the simplified example of a reduced two-species model in Section 4.4, that such a limit to “instantaneous” boundary conditions does not appear to be feasible in our modeling context. Beyond this apparent mathematical necessity, there appears to be biological data suggesting the presence of such regions close to the body edges, which separate head- and tail regions from the trunk, and the concept of pole regions is thought to play a fundamental role in regeneration, see [64].

Numerical discretization provides yet a different way to rationalize these dynamic boundary conditions. Representing the main body by grid points *U*_1_,…, *U_N_*, each carrying mass *hU_j_* where *h* is the grid size, we would simply add extra grid points *U*_0_ = *U*_−_ and *U*_*N*+1_ = *U*_+_, which carry mass *γU_j_, j* = 0, *N* +1. This can now be thought of as an inhomogeneous spatial grid, imposing no-flux boundary conditions on the thus extended domain. Clearly, letting the size *γ* of the boundary compartment decrease to the order of the grid scale *h*, we loose the concept of dynamic boundary conditions and should interpret the additional reaction terms 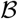 as nonlinear fluxes. We are not aware of a systematic analysis of such limits, connecting discretization, nonlinear fluxes, and dynamic boundary conditions.

#### Sensitivity to *wnt*-signaling gradients

The additional terms 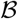 in the boundary dynamics model the strong local peak of differentiation of stem cells at wounds inside a formed blastema. In our model, this occurs in a region at the end of the interval considered. The differentiation process appears to be guided by *wnt*-signaling [2] and preserves polarity. Cutting experiments, where body fragments from quite different regions of a planarian regenerate while preserving polarity suggest that absolute values of *wnt*-signaling do not provide the relevant information. In light of these experiments, and in order to restore polarity from thin fragments regardless of specific location of the fragment inside the animal, it therefore seems necessary that stem cells detect gradients of *wnt*-related signaling and differentiate accordingly. Indeed, inspecting the *normal derivative* of *w* at the boundary after cutting in Figure 2.1, one readily sees that the sign of the normal derivative gives clues as to whether differentiation towards head or tail cells should occur.

Sensing of the gradient of *wnt*-signaling can be accomplished by stem cells in many ways, for instance during their (directed) movement towards the wounding site. During their movement they may measure the signal at different time steps or different locations, see [5], [26]. We do not attempt to model details of this sensing process and simply include a lumped reaction term for the differentiation of stem cells that depends sharply on the sign of the normal derivative of *w*.

Specifically, we include proliferation and terms for differentiation into the dynamics of *s*_±_, *h*_±_, and *d*_±_, of the form

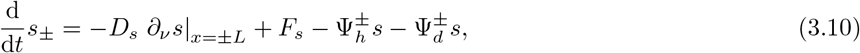

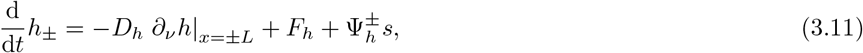

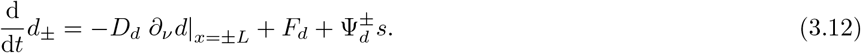

The functions 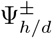 select differentiation of stem cells specifically in the boundary compartments, triggered by the lack of either head or tail cells, and directed according to the sign of *∂_ν_w*,

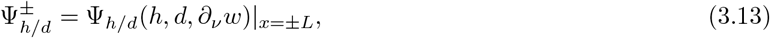

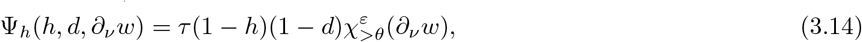

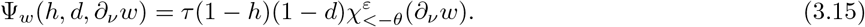

Here, *τ* is the rate of this mechanism and 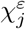 are smooth versions of the characteristic function given in (1.1); see Figure 3.1 for a schematic representation of the indicator functions *χ*.

### 3.3 Cutting, grafting, and growth

We present numerical simulations of our mathematical model illustrating homeostasis, cutting, grafting, and growth, confirming and expanding on the schematic representation in Figure 2.1.

#### Parameter values: diffusivities and rate constants

We chose parameter values representing roughly expected orders of magnitude within the system. Moderate changes of parameters do not have a significant effect on the outcome of simulations but we discuss some notable exceptions, below. The following table shows default parameter values, used unless noted otherwise. Random motion of stem cells is of order one as they move fairly freely through the body. Recall that we do not model directed motion of stem cells but recognize that such mechanisms may be necessary to accomplish gradient sensing in a robust fashion. Random motion of head and tail cells is very slow. Similarly, local signals *u_h/d_* diffuse slowly, while the long-range *wnt*-related signal has a diffusion constant of order 1. We work on domains of length 10 and attribute a mass fraction *γ* = 0.3 to the boundary, unless otherwise indicated. We assume fast proliferation of stem cells *p_s_* ≫ 1 and fast signaling dynamics relative to cell differentiation, *r_j_* ≫ 1. Cell differentiation *p_h/d_* and apoptosis *η_h/d_* as well as differentiation at the tissue edges during wound healing *τ* occur on a time scale of order 1. The production rate of *wnt*-related signals *p_w_* ≫ 1 is comparatively fast. The constant *ε* is small, approximating discontinuous switching in the kinetics, the constant *θ* measuring minimal detectable strength of the gradient at the body/wound edge in *ε*-units is set to 3. We comment below on some of the effects of changing parameter values but did not attempt a systematic study of parameter space.

#### Numerical implementation

We implemented the dynamic boundary conditions as time-dependent Dirichlet conditions. We then solved the system with grid spacing *dx* = 0.01 and time stepping 5 · 10^−4^ using a semi-implicit Euler method. We found little changes from refining spatio-temporal grids and also used Matlab’s stiff solver ODEİ5s for comparison with good agreement.

#### Homeostasis

We obtained equilibrium profiles starting with initial conditions that represent head and tail cells in the boundary compartments at the body edges, a uniform distribution of stem cells throughout the trunk, and a uniform *wnt*-signaling gradient. Solving the initial-value problem for a short time until, we found that concentrations approached constants in time.

Specifically, we used initial conditions

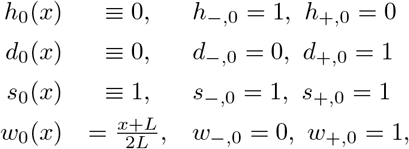

and let *u_h_* and *u_d_* equal *h* and *d*, respectively, for the initial data. The results, schematically shown in Figure 2.1, are displayed in Figure 3.2 and illustrate the stationary profiles of healthy organisms of different body size. We found that the linear concentration profile in the *wnt*-signal is quite robust under dramatic changes in the domain size. Fixing the width of the boundary compartment *γ* and varying *L* we found robust homeostasis between *L* = 0.005 (!) and *L* = 40. For very small L, the total variation of the *wnt*-signal *w* decreases: *w* stays bounded away from 0 and 1, and the gradient of *w* remains bounded as *L* tends to zero. In this regime, the signals *u_h_* and *u_d_* follow the concentrations *h* and *d*, respectively, less closely, being much smaller in amplitude. On the other hand, for very large *L*, the equilibrium state is sensitive to small fluctuations since the *wnt*-signaling gradient is small, within the interval [−*θ · ε, θ · ε*] where detection of the gradient is no longer decisive. Increasing the size of the boundary compartment *γ* helped stabilize dynamics generally and we found robust homeostasis for domain sizes 1,…, 50 and boundary compartments of size *γ* = 0.1,…, 15. Very small sizes of the boundary compartment, such as *γ* = 0.015, *L* = 5, were not able to sustain a head-trunk-tail profile, consistent with our discussion in Section 4, below. Whenever we saw homeostasis, we also tested robustness against small amplitude perturbations and found recovery within expected limits. A typical dangerous perturbation would for instance attempt to alter the sign of the *wnt*-signaling gradient near the boundary. We discuss perturbations more in line with some experiments, below.

**Figure 3.2:**
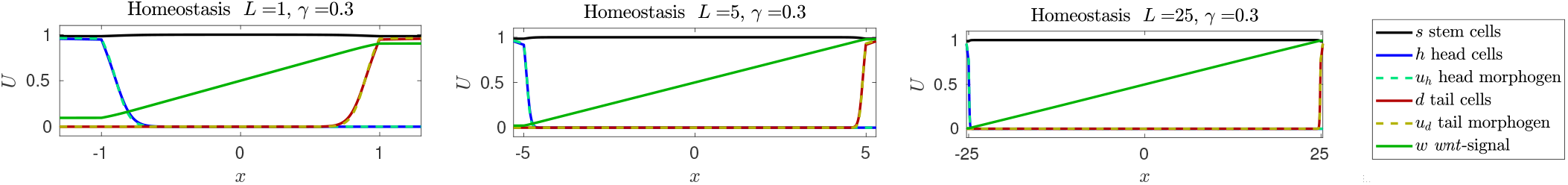
Equilibrium profiles showing a linear *wnt*-signaling profile, head and tail cells concentrated near boundaries, stem cell concentrations with small deviations from constant, and chemical signals closely following head and tail-cell concentrations, respectively. Note the different scales for *x*, which represent different body sizes of planarians. Homeostasis and subsequent cutting experiments are illustrated in the supplementary materials cutting_sequel.mp4.

The localization of the regions occupied by head and tail cells depends first on the strength of random motion *D_d_, D_h_* of *d* and *h* and diffusion rates of their associated morphogens; see Figure 3.3. On the other hand, changing production and degradation rates *r*_0_, *r*_1_, *r*_2_, *r*_3_ for *u_h/d_*, one can trigger a spontaneous expansion of the region of head and tail cells, where now the rate of expansion depends on these rates and the diffusivities.

**Figure 3.3:**
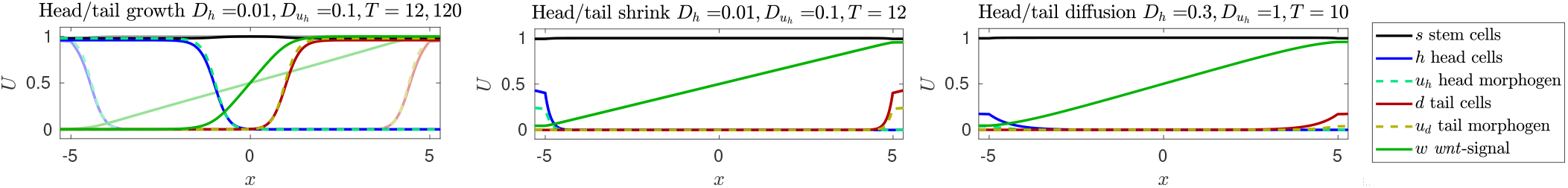
Growing and shrinking of head and tail cell regions (left/center) with *r*_0_ = 16, *r*_3_ = 4 (left) and *r*_0_ = 18, *r*_3_ = 10 (center); initial conditions: *s* ≡ 1, linear *w*-gradients, no head or tail cells. Expansion of head/tail regions shown on the left with snapshots at *T* =12 (opaque) and *T* = 120 (solid); shrinking terminates and homeostasis is reached at *T* = 12. Right, influence of strong random motion and diffusion on homeostasis. Throughout, parameters are as in Table 1 unless noted otherwise. Compare also the visualization in the supplementary materials cutting_sequel.mp4.

#### Cutting

Relating to experiments where a strip of the trunk of a planarian is cut out, we now study initial conditions that arise when using part of the homeostatic state in the central region of the domain, that is,

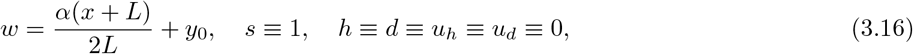

where *α* represents the fraction of the fragment cut out, and *y*_0_ ∈ [0,1] the *w*-concentration at the left edge of the cut fragment, corresponding to a cutting location *x*_0_ = *L*(2*y*_0_ − 1) ∈ [−*L, L*]. In other words, *α* = 0.02 corresponds to a cut of 2% of body length and *y*_0_ = 0.75 corresponds to a cut at three quarters of the body lengths distance from the head. Boundary data is chosen compatible *U*_±_ = *U*(±*L*) at time 0. Figure 3.4 demonstrates recovery from cuts of 2% and 4% of body length. Recovery is more sensitive if segments are cut out from tail or head regions. As expected, recovery depends on the sensitivity *ε*^−1^ and the threshold *θ*. Smaller values of *ε* and *θ* increase sensitivity and enhance recovery. On the other hand, small values of *θ* and *ε* also enhance vulnerability to noise and its effect on *wnt*-signaling gradients near the boundary. Smaller mass fractions *γ* ≲ 0.03 in the boundary compartments for a domain with *L* = 5 prevent recovery even for medium-size cuts, to the same extent as homeostasis breaks down. Generally, for cuts near the tail, the head regenerates first, and vice-versa. Throughout the simulations, polarity is consistently preserved.

**Figure 3.4:**
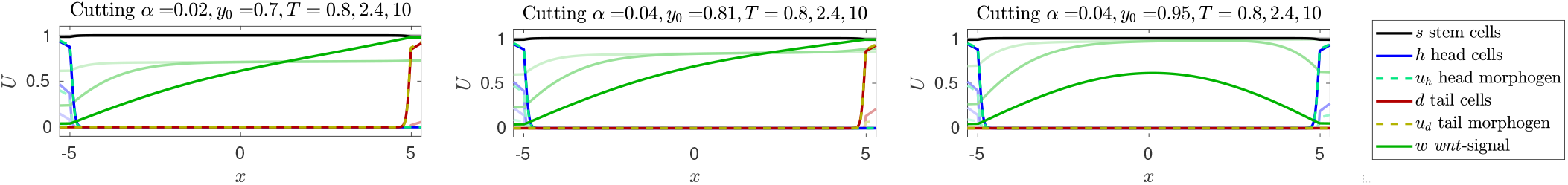
Regeneration after cutting experiments: cutting 1/50th from trunk region *y*_0_ = 0.7 (left), 1/25th close to tail *y*_0_ = 0.815 (center) and from tail *y* = 0.95 (right); see (3.16) for initial conditions. Note the failure to regenerate through the emergence of heads on both sides and the loss of the *wnt*-signaling gradient on the right, for a cut very close to the tail region. Throughout, parameters are as in Table 1; opaque curves show concentrations at earlier snapshots as listed in the title.

In our numerical simulations, one observes an initial strong burst in differentiation triggered by the boundary sensing 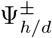. This is compensated for by the strong proliferation of stem cells *p_s_*. Simulations still show a resulting decrease in the stem cell population at the boundary, 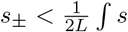, which can be much more pronounced for different parameter values such as smaller *p_s_* or larger *τ*. This reflects and quantifies in our model the experimentally observed strong proliferation of stem cells during wound healing.

#### Grafting

In this third set of numerical experiments, we study initial conditions where head cells are inserted into a healthy planarian. We therefore use the results of the homeostasis simulation as initial condition and then change values in a region of size 10% of the length of the organism. In this region, we increase the concentration of head cells *h* and the associated morphogen *u_h_* to 1 and eliminate *wnt*-signaling, that is, set *w* ≡ 0. We observe that the head region survives and changes the profile of *wnt*-signal concentrations. Grafting head cells of a donor close to the head region of the host leads to merging of the two head regions. Grafting donor head cells near the tail of the host can preserve the tail, unless tail cells are significantly destroyed during grafting. Figure 3.5 shows numerical experiments corresponding to these three scenarios. We think of these different outcomes, for instance the possibility of merging of heads vs persistence of the new head as reflecting the dichotomy seen in experiments where an additional head may grow out of a graft or the graft may just disappear; see the discussion in Section 2 and Figure 2.1.

**Figure 3.5:**
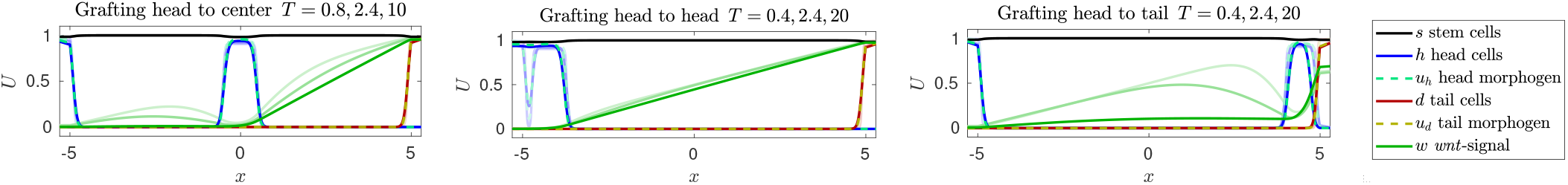
Regeneration after grafting head tissue into center of trunk (left), head region (center), and tail region (right). Throughout, parameters are as in Table 1; opaque curves show concentrations at earlier snapshots as listed in the title. Note in particular the persistence of the grafted head in the left and right panels versus the merging (and eventual vanishing) of the graft in the center panel. Compare also the visualization in the supplementary materials grafting.mp4.

We caution here that these numerical experiments rely to some extent on astute choices of production rates for head and tail cells and the associated morphogen, as well as the choice of diffusivity. In our experiments, the boundary between head and trunk cells is nearly stationary and sharply localized. Roughly speaking, an open set of parameter values will lead to expanding head (or tail) regions, a complementary open set will lead to shrinking head (or tail) regions. At the boundary of these parameter regions, head and tail regions will remain nearly stationary, subject only to a slow coarsening interaction as seen in the head-to-head grafting. Stronger diffusivities will enhance both speed of growth and shrinking as well as the slow coarsening. Our choices of parameters are close to the critical values, where head and tail regions neither shrink nor expand. We emphasize that for parameters where head- or tail regions shrink, grafted regions will slowly disappear, but the head- and tail regions near the boundary will persist due to the *wnt*-related gradient triggered differentiation of stem cells.

We did not explore the phenomena of growth and shrinking of planarians in much more detail than in the next paragraph, since our model so far does not include any well-motivated mechanisms that would regulate the size of organs.

We note that regions between two heads (or between two tails) do not generate a *wnt*-signaling gradient. One would therefore predict that secondary cutting experiments with tissue from such regions after grafting would not be able to regenerate consistently and not preserve polarity.

Mimicking dissociation experiments in hydra, we also explored outcomes of random grafting in our mathematical model, see Figure 3.6. We inserted head and tail fragments of roughly 3% of body length randomly at various locations in the simulated planarian. Whenever these pieces are large enough, they persist as head and tail regions; smaller regions eventually disappear.

**Figure 3.6:**
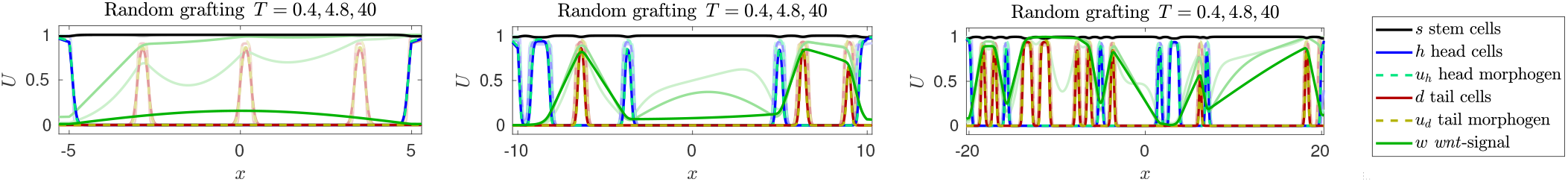
Regeneration after grafting multiple head and tail regions, for different body lengths *L* = 10, 20, 40. Throughout, parameters are as in Table 1; opaque curves show concentrations at earlier snapshots as listed in the title. Annihilation of all grafts on the left, and multiple persistent regions for larger domains (center and right).

#### Growth

As a last experiment, we demonstrate that the mechanisms of our model are capable of sustaining robust patterning when the body size of the planarian expands. In our simulations, the body size expands uniformly at a constant speed *c* = 0.3, 1.5, 3 and we monitor the concentration profiles; see Figure 3.7. More precisely, we assume that tissue locations evolve according 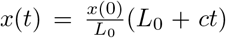, where (−*L*_0_, *L*_0_) is the body size at time *t* = 0 and (−*L*(*t*), *L*(*t*)), *L*(*t*) = *L*_0_ + *ct*, is the body size at time *t*. Mass conservation in the extended domain, in the absence of reaction terms, then forces the dilution *∂_t_U*(*t,x*) = *ρ*(*t*)*U*(*t,x*), with 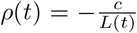. Transforming back to a fixed domain with a coordinate change *x* ↦ *xL*_0_/*L*(*t*) gives a diffusion equation on (−*L, L*) with diffusion matrix 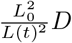. In the abstract formulation (3.1), we therefore simply amend the diffusion constant in our model and add a dilution term to model the system on the growing domain,

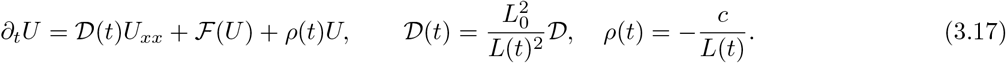

**Figure 3.7:**
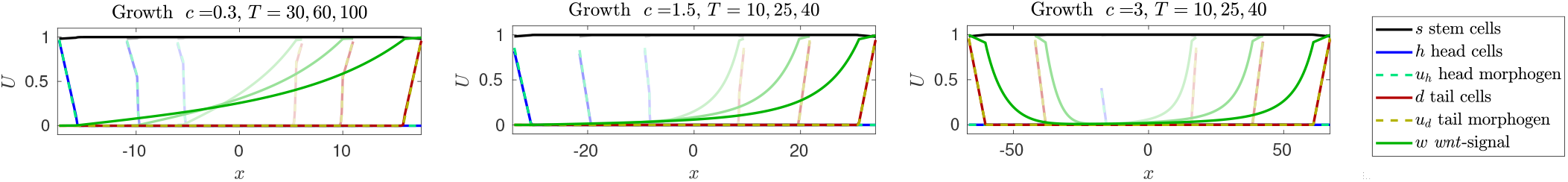
A healthy planarian under uniform linear growth with speeds *c* = 0.3, 1.5, 3; see text for more details. Concentration profiles are plotted in actual coordinates, such that they occupy only part of the final domain at earlier times (faded curves terminate at ±(*L*_0_ + *cT*)). Profiles are well maintained close to homeostasis for slow (left) and moderate (center) growth speeds. For larger speeds (right) dilution of *w* reduces the overall concentration such that the gradient at the left boundary where *w* ∼ 0 falls below the sensing threshold. As a result concentration of head cells decreases on the left boundary and a second tail replaces the head.

We chose to preserve the relative size of the boundary compartments such that 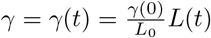, which introduces the same dilution term in the equation for the dynamic boundary values *U*_±_. Fluxes need to be adjusted to the changing diffusion constant 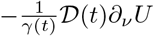. We refer to [76] for a more detailed discussion of different growth laws and references to the literature.

We simulated the resulting system on a fixed grid corresponding to scaled *x*-coordinates but plotted the outcome in the actual unscaled *x*-variables. Parameters were chosen according to Table 1. The results are displayed in Figure 3.7. For slow and moderate growth speeds, the concentration profiles resemble the homeostatic profiles at a given length. We observe failure to maintain key features only for quite rapid expansion (the time scales of *c* = 3 corresponds to a doubling of a planarian of size 9 in 3 time units, while cell differentiation happens with rate 1, generating at most 1 unit volume of head or tail cells from stem cells in one time unit). This failure at rapid growth can be attributed to the dilution of the *wnt*-related signal that is not adequately compensated for by production through tail cells and diffusion. As a result, the *wnt*-signaling gradient becomes very small in the head region. As already demonstrated in the case of fixed domains during homeostasis and cutting, solutions become sensitive to perturbations when *wnt*-signaling gradients approach the sensing limits determined by *ε* and *θ*.

**Table 1:**
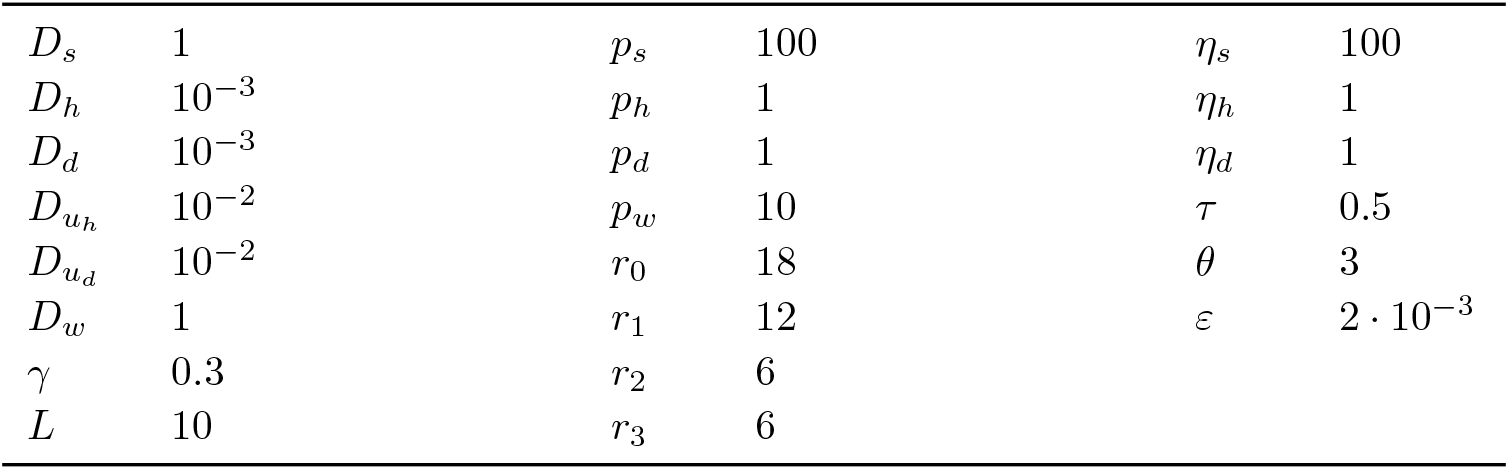
Parameter values used in the numerical simulations throughout, unless noted otherwise.

|  |  |  |  |  |  |
| --- | --- | --- | --- | --- | --- |
| $D_s$ | 1 | $p_s$ | 100 | $\eta_s$ | 100 |
| $D_h$ | $10^{-3}$ | $p_h$ | 1 | $\eta_h$ | 1 |
| $D_d$ | $10^{-3}$ | $p_d$ | 1 | $\eta_d$ | 1 |
| $D_{u_h}$ | $10^{-2}$ | $p_w$ | 10 | $\tau$ | 0.5 |
| $D_{u_d}$ | $10^{-2}$ | $r_0$ | 18 | $\theta$ | 3 |
| $D_w$ | 1 | $r_1$ | 12 | $\varepsilon$ | $2 \cdot 10^{-3}$ |
| $\gamma$ | 0.3 | $r_2$ | 6 | | |
| $L$ | 10 | $r_3$ | 6 | | |

Of course, the externally imposed growth used here is not a completely appropriate representation of the actual growth of planarians while feeding. Conversely, the model can not yet clearly characterize the dynamics under starvation conditions when sizes of planarians shrink by an order of magnitude: the negative dilution term would give an overcrowding of cells with sometimes unstable signaling gradients near the boundary. A more accurate model would relate growth laws and proliferation, which in turn depends on food supply. Nevertheless, the scenario studied here does show the inherent independence of the patterning from body size and its robustness for growing planarians.

## 4 Analysis via model reduction

We first argue that stem cells and short-range signals can be eliminated in a quasi-steady state approximation in Section 4.1. We reduce the resulting system for (*h, d, w*) further by considering one order parameter *c* for the cell type instead of the two concentrations (*h, d*) in Section 4.2 and illustrate how this reduced two-species system captures regeneration after cutting and grafting in Section 4.3. Finally, we focus on the key element of the process, the restoration of the *wnt*-related signal in a further reduced scalar equation.

### 4.1 Eliminating stem cells and short-range signals

A simple analysis of kinetics, supported by observations in numerical simulations, suggests two quick simplifications by reduction.

#### Stem cell populations are constant in time and space

Inspecting the proliferation and differentiation rates of stem cells, one notices that proliferation is much faster than differentiation and equilibrium concentrations depend only marginally on differentiation, that is, on the concentrations of *u_h/d_*. Setting *p_s_*(*s*) = 0, we then obtain equilibrium concentrations *s* = (*p_s_* − *η_s_*)/*η_s_*, namely, in our choice of parameters, *s* ≡ 1.

#### Short-range signals are tied to cell populations

Assuming in the following that *s* ≡ 1, we notice that the signal production rates *r_j_* are much larger than the cell differentiation and death rates *p_h/d_* and *η_h/d_*. Using an adiabatic reduction for the kinetics, one then equilibrates the reaction rates for *u_h/d_* and finds *u_h/d_* as functions of *h* and *d*,

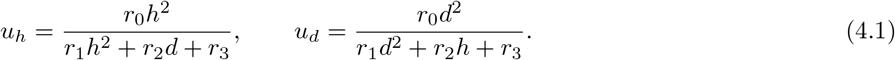

Similar to the case of stem cell dynamics, the reduction for the kinetics is valid under suitable bounds on gradients, and we shall demonstrate below that such reduced systems capture the key structure of the dynamics quite well.

In summary, the reduction thus far, substituting the expressions for *u_h/d_* into the equations for *h, d, w*, after setting *s* ≡ 1, yields the system

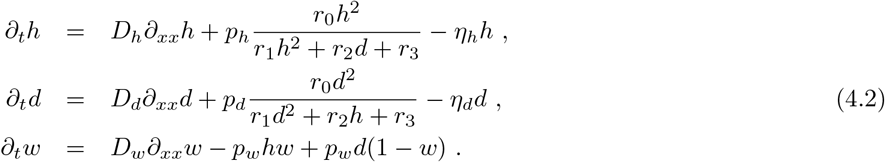

For the boundary data, we find

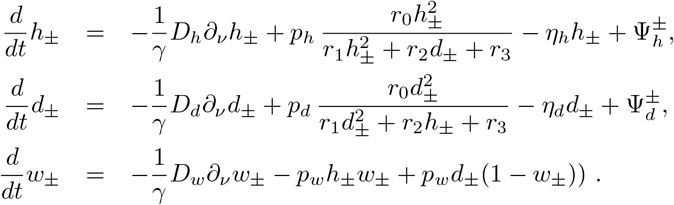

where again

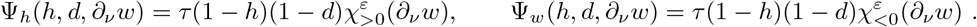

Simulations of this reduced model are almost indistinguishable from the full model and therefore we do not display the somewhat redundant graphics, here. We proceed by a further, more phenomenological simplification.

### 4.2 From cell type to order parameter

In numerical simulations of our mathematical model, which are related to the main biological experiments described above, concentration profiles of *h* and *d* are mostly constant, taking on values (*h*_*_, 0), (0, *d*_*_), or (0, 0), where in fact *h*_*_ = *d*_*_ = *h*_+_,

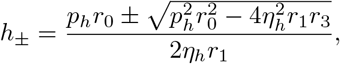

provided that 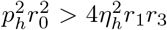, which we shall assume in the sequel. These three states are in fact stable equilibria for the ODE kinetics and for the full PDE (4.2) for *h, d*, (due to the equal diffusivites) and correspond to head-only, tail-only, and trunk-only states, respectively. The equilibria (*h*_−_, 0) and (0, *h*_−_) satisfy 0 < *h*_−_ < *h*_+_ and are unstable threshold states, separating initial conditions that evolve toward head cells from initial conditionsevolving toward trunk-only cells. Due to the symmetric choice of parameter values, pure head and tail states have equal concentration values here, although this is not necessary in order to capture the considered phenomena.

Regions of constant values of *h* and *d* are separated by interfaces (or fronts) that propagate with a speed determined by the parameter values *r_j_*, *η_j_*, and *p_j_*. There are three possible fronts, head-trunk, tail-trunk, and head-tail.

Fronts between pure head and trunk state can be understood in the *h*-equation alone, setting *d* ≡ 0. This equation

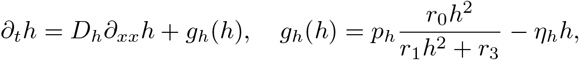

is in fact a gradient flow related to the free energy 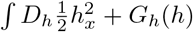. Here 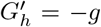 is the potential with critical points 0, *h*_±_. The direction of motion of the front changes sign at the Maxwell point, when *G*(*h*_+_) = *G*(0). For the parameter values chosen in our simulations, fronts propagate slowly toward the head region. Fronts between tail and trunk regions are completely equivalent. Fronts between tail and head regions do not propagate if they exist. Depending on parameter values, such fronts may not exist but front-like initial conditions rather split into two fronts (*h*_*_, 0) ↔ (0, 0) ↔ (0, *h*_*_), where the newly emerged state (0, 0) expands. All of those features can be captured in a scalar equation. Somewhat formally, on the level of the kinetics, one can envision straightening out the line segments in the *h* − *d*-plane (*h*_*_, 0) → (0, 0) → (0, *d*_*_) into a one-dimensional line segment −1 → 0 → 1; see Figure 4.1. One then arrives at the scalar equation

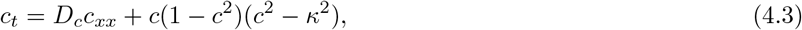

with 0 < *κ* < 1. Equation (4.3) possesses stable equilibria {−1, 0, 1} corresponding to (1, 0), (0, 0), and (0,1), respectively, and possesses reflection symmetry between *c* → −*c*, mimicking the reflection symmetry between *d* and *h* in our simplified model. Here, the critical “Mawell” point is 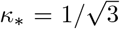, such that (0, 0) invades (1, 0) and (0,1) for *κ* < *κ*_*_ and (0, 0) is invaded otherwise. Similarly to the systems case (4.2), fronts between −1 and 1 alias head and tail exist in (4.3) precisely when splitting of the front is not expected, that is, when −1 invades 0. This can be readily seen from the phase portrait of the steady-state equation.

**Figure 4.1:**
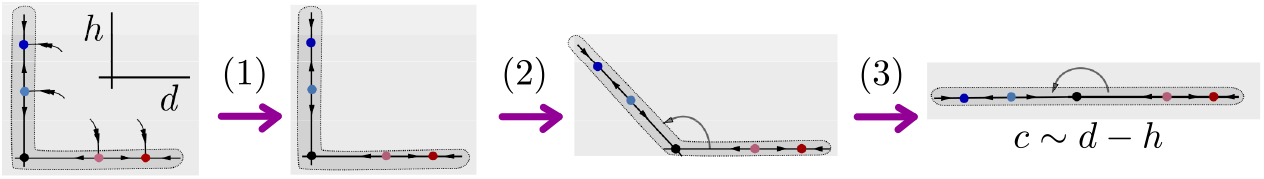
Schematic of the reduction from cell type to order parameter. In the kinetics of the head-cell dynamics, the *ω*-limit set of a large ball consists of the line segments of the coordinate axis between the origin and pure-head and pure-tail state, respectively. Restricting to this *ω*-limit set (1) gives the joined line segments, which one then (2) bends open to (3) arrive at a straight one-dimensional line segment with dynamics equivalent to the scalar order parameter *c* ∼ *d* − *h*; see text for a rationale for this reduction including diffusive effects, particularly fronts.

The dynamics for *wnt*-related signal concentrations need to be modified accordingly, with production and degradation now occurring for *c* positive and negative, respectively, for instance through a term

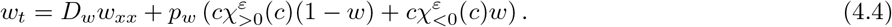

Gradient sensing at the body edge is lumped together to yield a production term in the equations for the boundary concentrations *c*_±_ of the form

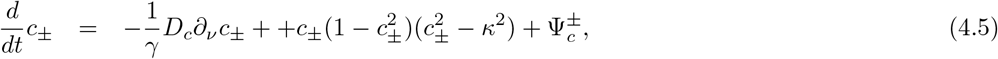

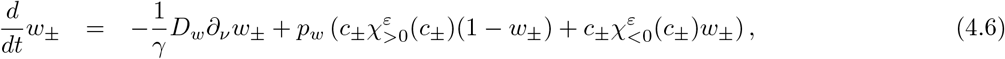

where

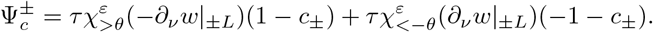

The set of equations (4.3)–(4.4) with dynamic boundary conditions (4.5)–(4.6) form the minimal reduced system that is able to mimic robust regeneration under cutting, grafting, and growth, which we shall demonstrate in the next section.

### 4.3 Homeostasis, cutting, and grafting in the reduced model

We simulate system (4.3)–(4.6) with parameter values

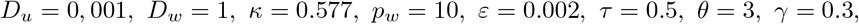

on a domain of size 2*L* = 10. Figure 4.2 illustrates the prototypical experiments and regeneration with these parameter values. Initial conditions for cutting are

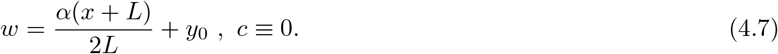

**Figure 4.2:**
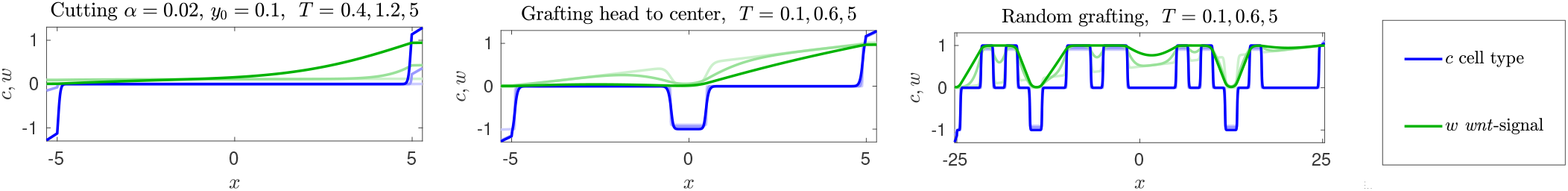
Simulations of the reduced model (4.3)–(4.6) with initial conditions corresponding to a cutting experiment (left), a grafting experiment where a head is grafted into the center portion (middle), and a random grafting of head and tail pieces (right); see text for details on initial conditions.

Initial conditions for grafting are the homeostatic equilibrium with *w* = 0 and *c* = ±1 on segments of length 1.

Note that we chose *κ* near the Maxwell point, which prevents changes of size in grafting experiments. Different choices lead to expanding or retracting head- or tail regions, as shown in the full model in Figure 3.3.

Increasing *ε* or decreasing the mass in the body edge compartments *γ* significantly will prevent recovery. Near critical values, we observe non-monotone behavior of the gradient and the boundary values *w*_±_; see Figure 4.3. We will point to some explanation in the next section when we discuss an even further reduced, scalar model. As in the full model, we see a dichotomy for head grafts, between merging of the two heads for grafts near the head of the host, and persistence of one head only for grafts sufficiently far away from the head of the host. This we relate to the dichotomy between vanishing of the graft and outgrowth of a new head in experiments.

**Figure 4.3:**
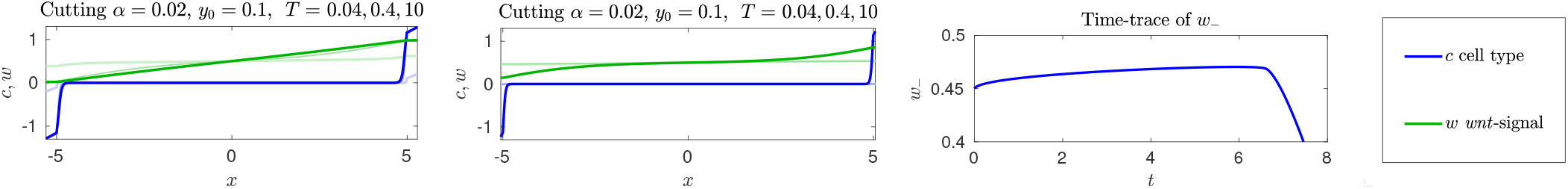
Simulations of the reduced model (4.3)–(4.6) with initial conditions corresponding to cutting experiments, with parameter choices causing difficult recovery or recovery failure. Larger *ε* = 0.012 leads to slow and non-monotone recovery (left; associated time series of *w*_−_ on the right): the *wnt*-signaling gradient first decreases before head and tail cell populations are established. The same phenomenon occurs when reducing the mass in the boundary compartment *γ* = 0.055 while keeping *ε* = 0.002 (center). Larger values of *ε* and smaller values of *γ* typically prevent recovery.

### 4.4 Analysis and comparison with Robin boundary conditions

Throughout this section, we restrict to sensitivity thresholds *θ* = 0. System (4.3)–(4.4) allows for a trivial solution *c* ≡ 0, *w* ≡ 1/2. In fact, cutting experiments with small fragments can be thought of as starting with initial conditions close to this trivial solution. Regeneration and regeneration failure from small fragments can therefore be well predicted from a stability analysis of this trivial solution: stability would prevent regeneration from small fragments, while instability indicates the initial phase of the pattern-forming process.

Linearizing at the trivial solution, we find the system

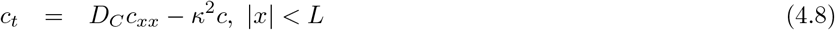

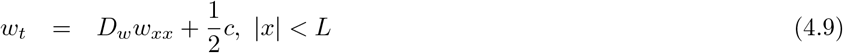

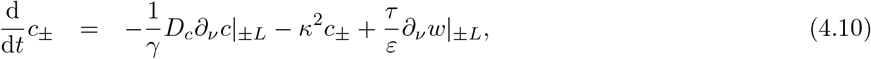

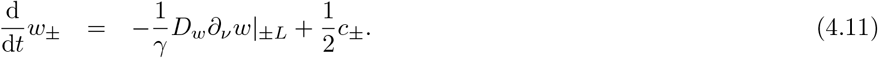

After Laplace transform, the associated eigenvalue problem can be converted into a transcendental equation which however is difficult to analyze theoretically. Numerically, one readily finds instabilities in the relevant parameter regimes. We illustrate here the basic instability mechanism with some formal simplifications.

Consider the left boundary, such that the eigenfunctions are of the form 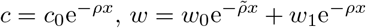, where *ρ* and 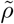 are determined by the spectral parameter λ, 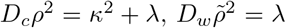, and *w*_1_ depends on *c*_0_. Substituting these expressions into the equations on the boundary yields a generalized eigenvalue problem.

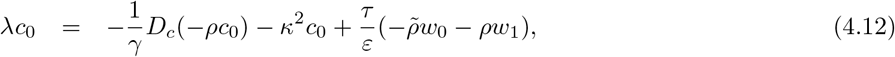

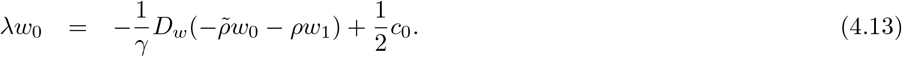

The key ingredient here are the off-diagonal terms: 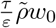 from 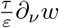 in the equation for *c*_0_, and 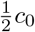 in the equation for *w*_0_. For positive λ, these terms are both positive hence causing a negative determinant and hence instability.

These off-diagonal terms can also be understood more directly but less quantitatively as encoding a positive feedback mechanism. A small increase of *w*_−_ will cause a positive normal derivative of *w* and hence an increase in *c*_−_ at the boundary from the equation for *c*_−_. The increase in *c*-translates into a further increase of *w*_−_ through the term 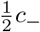 in the equation for *w*_−_, thus providing the positive feedback mechanism responsible for instability.

It is here that we can see how dynamic boundary conditions are a key ingredient. Relaxing these dynamic boundary conditions by letting reaction rates at the boundary tend to infinity, we formally obtain the mixed Robin boundary conditions

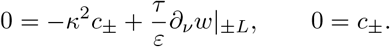

One readily sees that, for well-posedness, the condition from the equation for *c*_−_ needs to be associated with the equation for *w* since it contains the *w* normal derivative, while the condition from the second equation for *c*_−_ simply gives a Dirichlet condition for the *c*-equation. As a consequence, the *c*-equation decouples as a simple diffusion equation with decay −*κ*^2^*c*_−_ and Dirichlet boundary conditions. Setting *c* = 0 then gives a diffusion equation for *w* with homogeneous Neumann boundary conditions, which implies stability.

More directly, this calculation points to the failure of a much simpler model. Indeed, we could model an instantaneous adaptation of tail and head cell concentrations, alias the order parameter *c*, to the sign of the signal gradient through *c* = sign(*∂_ν_w*), with a possibly smoothed out version of the sign-function. One would then let *w* follow head and tail cell concentrations encoded in *c*, for instance through *w* = *c*. This approach is bound to fail, since the boundary condition *c* = sign(*∂_ν_w*) will *not* enforce *c* to increase from 0 to 1, say, in a cutting experiment where *∂_ν_w* > 0. Instead, the boundary condition can be satisfied by an, in comparison, much smaller adjustment of the *w*-levels near the boundary which achieves *∂_ν_w* = 0, lowering for instance the concentration of *w* only slightly in a region close to the boundary. The value *∂_ν_w* = 0 then is compatible with *c* = 0 in the boundary condition (with the convention sign(0) = 0). We tested such boundary conditions numerically and observed a subsequent decay of the *w*-gradient and convergence to *c* = 0, that is, failure of regeneration. Relating to the previous discussion, the boundary condition *c* = sign(*∂_ν_w*) fails to enforce regeneration since it is not explicitly associated with a forced change in levels of *c*, but acts rather as a nonlinear flux for the *w*-equation. Dynamic boundary conditions provide precisely this association and therefore guarantee regeneration.

We numerically confirmed these observations on the necessity of dynamic boundary conditions. We observed stability both in a discretization with mixed boundary conditions and when reducing mass fractions to the scale of the grid size.

### 4.5 Recovery of the long-range signal gradient as organizing feature

We can achieve a further simplification to a single scalar equation if we focus on experiments with only head and tail at the respective extremity, excluding in particular grafting experiments, and tracking only *wnt*-related concentrations. We assume that the order parameter *c* mostly vanishes in the domain, that is, head- and tail-cells are confined to the boundary regions. In this case, the kinetics for *w* vanish inside the bulk |*x*| < *L*, and we are left with a simple diffusion equation for *w*. Note that his explicitly excludes grafting experiments and more generally states with multiple head and tail regions.

With this assumption it is also sufficient to track the boundary data *c*_±_ for the order parameter. Assuming now that *c*_±_ adjusts rapidly according to (4.5), we can set *c*_±_ ∼ sign *∂_ν_w*. This all gives us the simple scalar equation

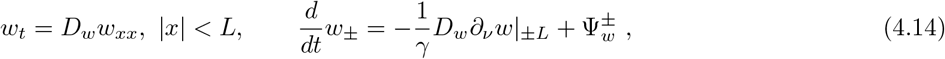

with

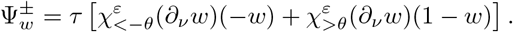

#### Cutting — numerical experiments

We first illustrate regeneration of the *wnt*-signaling gradient after cutting, and therefore use parameters

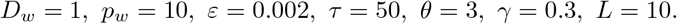

Figure 4.4 shows recovery from small fragments cut from center- or head regions. Recovery is slightly less robust for fragments cut near head or tail. For increasing values of *ε* which amounts to decreasing the sensitivity of the gradient sensing at the boundary (or, also, for increasing values off the mass fraction *γ*), we see a transition to a system state which fails to recover; see Figure 4.5. One first sees oscillatory decay toward the trivial state *w* ≡ 0.5 before sustained oscillations emerge in a weakly subcritical Hopf bifurcation. The unstable periodic orbit separates initial conditions that lead to the trivial, trunk-only state *w* ≡ 0.5 from initial conditions that converge to large-scale oscillations. The large oscillations eventually appear to terminate in a heteroclinic bifurcation for larger values of *ε*. We saw qualitatively similar transitions when decreasing the mass fraction *γ* instead of increasing *ε*.

**Figure 4.4:**
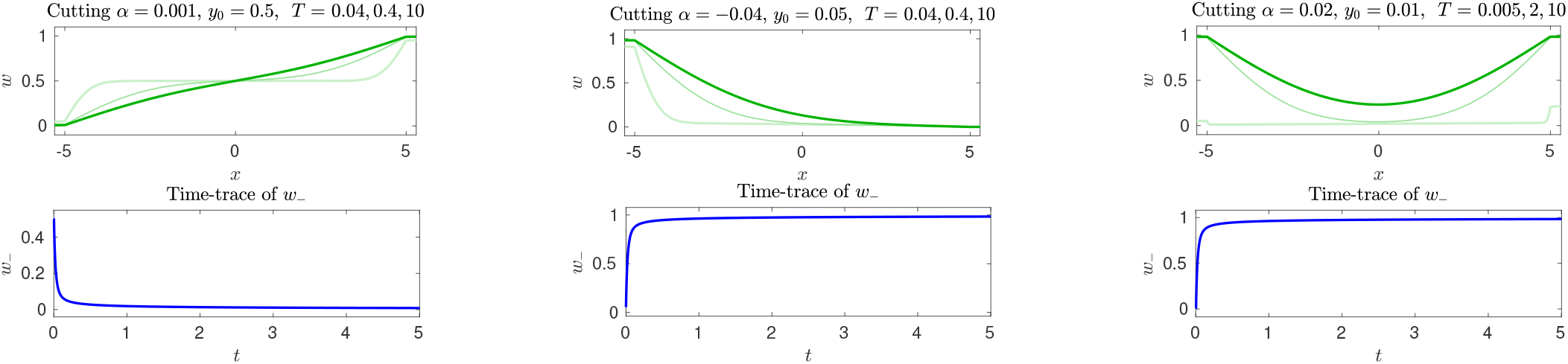
Simulations of the scalar model (4.14) with initial conditions corresponding to cutting experiments, that is, linear profiles in *w* with small slopes. The left panel shows robust recovery for a very small fraction, 0.1%, from the center region. The center panel shows recovery for a somewhat larger fraction, 4%, from the head region. Here the cut is reflected to demonstrate that recovery is independent of the orientation of the *x*-interval. The right panel shows failure of recovery when a slightly smaller fragment is cut from the head region, in agreement with experimental observations that recovery from fragments near head and tail is less robust; see text for details. Note the slightly different time instances shown in the right panel, illustrating the very rapid failure to preserve the sign of *∂_ν_w* near the boundary. The second row shows corresponding time series of *w*_−_.

**Figure 4.5:**
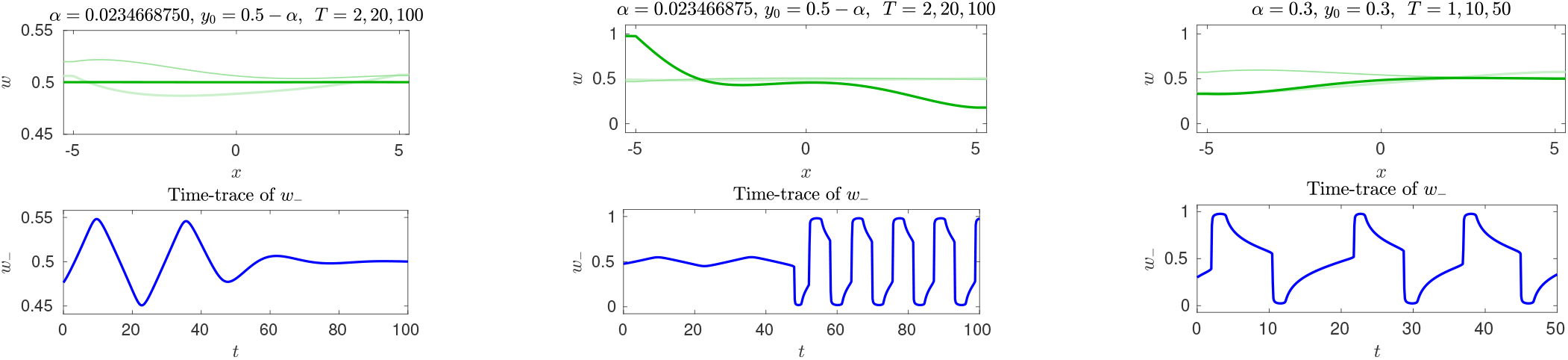
Simulations of the scalar model (4.14) with initial conditions corresponding to a cutting experiments and moderate value of sensitivity *ε*^−1^ = 15. An unstable oscillation separates initial conditions w.r.t. *α* that decay to *w* = 0.5 (left) from initial conditions with sustained large-amplitude oscillations (center). Note the different scales for *w* here. Oscillations disappear in a saddle-node of periodic orbits for smaller values of *ε*. For yet smaller sensitivity *ε* ∼ 12.5, the oscillations seem to disappear in a heteroclinic bifurcation (right). Corresponding time series of *w*_−_ in the second row. See text for predictions of Hopf bifurcations and implications for model corroborations. Oscillations are illustrated in the supplementary materials scalar_oscillations.mp4.

#### Analysis — equilibria and stability

We assume that *τ* ≫ 1 and neglect the flux term 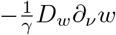. We also assume that *θ* = 0 such that the characteristic function takes value 1/2 at *w* = 1/2. At the boundary, equilibria then satisfy either,

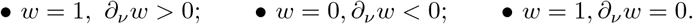

For moderately sized domains, *L* ≪ *ε*, this yields three equilibria, *w* ≡ 1/2, *w* = (*x*+*L*)/(2*L*), and *w* = (−*x*+*L*)/(2*L*), corresponding to only trunk, head-tail, and tail-head solutions. Linearizing at these equilibria, we find a diffusion equation with dynamic boundary conditions

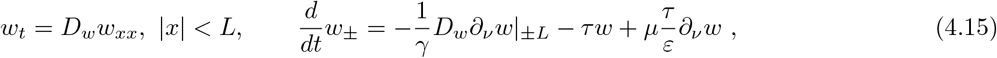

where *μ* = 1 for the trunk solution *w* ≡ 0 and *μ* = 0 for the head-tail and tail-head solutions.

One readily finds that head-tail and tail-head solutions are always stable. To analyze stability of the trunk-only solution, *mu* = 1, we consider the semi-unbounded domain *x* > 0, set *w* = *e*^λ*t*+*ρx*^, with λ = *ρ*^2^, to find the characteristic equation

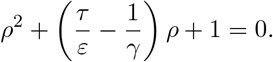

For *ρ* to correspond to an eigenvalue, we need Re*ρ* < 0, which implies 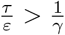. We find

- *real instability:* 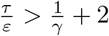 gives 2 real roots *ρ*_±_ < 0 and two associated real unstable eigenvalues λ_±_ > 0;
- *complex instability:* 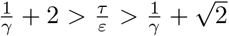 gives 2 complex conjugate roots with Re*ρ*_±_ < 0 and two associated complex conjugate unstable eigenvalues Reλ_±_ > 0;
- *Hopf bifurcation:* 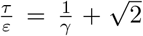 gives 2 complex conjugate roots with Re*ρ*_±_ < 0 and two purely imaginary eigenvalues Reλ_±_ = 0;
- *stability:* 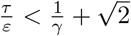 gives eigenvalues λ with negative real part.

We found that the Hopf bifurcation is subcritical, the unstable periodic solution stabilizes in a saddle-node bifurcation of periodic orbits, grows in amplitude and eventually disappears in a heteroclinic bifurcation for sufficiently large values of *γ*.

The dynamics are quite different if the boundary conditions are relaxed to Robin boundary conditions, for instance

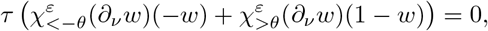

at *x* = ±*L*. Neglecting the flux term 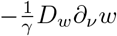, that is, with sufficiently large mass fractions *γ*, equilibria for dynamic and Robin boundary conditions coincide. Stability of equilibria is however quite different: the linearization with Robin boundary conditions is a Sturm-Liouville eigenvalue problem

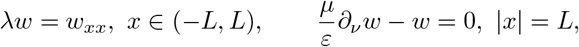

with real eigenvalues λ_0_ > 0 > λ_1_ > …, λ_0_ ∼ *ε*^2^/*μ*^2^ for *L* ≫ 1. The dynamics generally do not allow for oscillations as observed in the case of periodic boundary conditions. Nevertheless, the trivial trunk-only solution is unstable in this approximation and we see robust recovery of small *w*-gradients. One can attribute the appearance of oscillations to an effective distributed delay in the otherwise scalar equation for the boundary dynamics of *w*_−_ caused by the coupling to the diffusive field *w*(*t, x*) which acts as a buffer that stores a blurred history of boundary data.

The transitions discussed here occur when gradient dependence at the boundary is not sufficiently sensitive, that is, *ε* is not sufficiently small or *τ* is too small, or when the mass fraction in the boundary region *γ* is too small. This appears to be a prediction quite specific to this model, occurring to some extent also in the full system and the system with order parameter. We are not aware of experimental observations of oscillations in cases where recovery and regeneration are severely impeded. But any such observation would clearly corroborate our basic modeling assumptions.

## 5 Discussion

We presented a mathematical model for robust regeneration of planarians. Our model is able to reproduce most cutting and grafting experiments. As opposed to modeling efforts based on Turing-type mechanisms, our model preserves polarity after cutting and yields robust results over organism scales differing by factors of 100.

Central to our model are two observations from experiments:

i. sharply increased activity including stem cell proliferation near wound healing sites;
ii. global gradients of chemical signals, related to the *wnt*-signaling pathway.

We translate the first observation into dynamic boundary condition, modeling changed reaction kinetics in a boundary compartment. The second observation has often been discussed in connection with the regulation of tissue size, expanding on the idea of the French-flag model. The role of this global signal gradient is different in our model: we postulate that the gradient, rather than absolute levels of the signal are sensed by stem cell populations and, at wound sites, translated into directed differentiation. We suspect that within a rather general modeling context, such a gradient sensing is necessary in order to reproduce robust preservation of polarity in cutting experiments.

We incorporate these ingredients into a comprehensive model for 6 species, 3 cell type populations and 3 chemical signals. Through model reduction to an order parameter for cell types and one long-range chemical signal, only, we exhibit how these two ingredients organize the regeneration process. In the reduced model, we can point to regeneration as an instability mechanism for a trivial, unpatterened state and identify analytically limits of robust regeneration. The process is fundamentally different from Turing’s mechanism, and driven by the boundary compartments.

There are several ways in which the model could be refined. First, one can quite easily introduce a head-tail bias, which is clearly observed in experiments. This could be introduced either through different sensing thresholds *θ* at body edges, or through different differentiation and proliferation kinetics. One can thus introduce a bias towards head generation from small fluctuation of trivial states. We are not aware of a causal rather than a phenomenological justification of this bias. Second, we do not attempt to model regulation of the size of the head and tail regions. Since our model relies essentially on establishing a global signaling gradient, cells can clearly obtain positional information by reading out absolute levels of the *wnt*-related signal. Postulating such a dependence of differentiation or apoptosis on these levels, one can then introduce *w*-dependence in the {*s, h, d, u_h_, u_d_*}-subsystem and thus influence the tristability. Front motion as described in the reduced model in Section 4.2 would then depend on *w* levels, and stationary interfaces would lock into fixed *w*-levels, thus regulating a fixed size of head or tail as a percentage of the full body length. In the order parameter model, this could for instance be accomplished by a *w* dependence of the quintic kinetics,

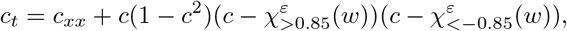

which would regulate the size of head and tail to regions where *w* > 0.8 and *w* < −0.8, respectively, about 15% of body size, each. Pushing this further, such effects could also model the spontaneous formation of head cell clusters when suppressing the *wnt*-signaling pathway. Lastly, one could address directed motion of stem cells or progenitors. Our focus here was on essential features enabling regeneration in a minimal model and we therefore ignored (active) transport other than random motion and diffusion. Since experiments appear to point to a significant role of directed motion, a model that takes such effects into account could potentially improve both qualitatively and quantitatively on the results presented here.

## Supporting information

Illustration of cutting as in Fig. 3.2 and 3.3

Illustration of oscillation as in Figure 4.5

Illustration of grafting as in Figure 3.5

## Acknowledgments

CT was supported by a fellowship of the Graduate School of the Cells-in-Motion Cluster of Excellence EXC 1003–CiM, University of Münster (WWU). ASc was partially supported through NSF grant DMS–1311740 and DMS–1907391, a Research Award from the Alexander-von-Humboldt Foundation, and a WWU Fellowship. ASc and ASt were supported by the DFG (German Research Foundation) under Germanys Excellence Strategy EXC 2044-390685587, Mathematics Münster: Dynamics - Geometry Structure. CT and ASt gratefully acknowledge numerous valuable discussions about the biology of planarians with Kerstin Bartscherer.

